# Cancer Interception During Treatment: Using Growth Kinetics to Create a Continuous Variable for Assessing Disease Response

**DOI:** 10.1101/2024.09.10.612311

**Authors:** Mengxi Zhou, Tito Fojo, Larry Schwartz, Susan E. Bates, Krastan B. Blagoev

**Affiliations:** Department of Medicine, Division of Hematology/Oncology, Columbia University Medical Center, New York, NY and James J Peters VA Medical Centre, Bronx, NY, USA; Department of Radiology, Memorial Sloan Kettering Cancer Center; National Science Foundation, Alexandria, VA 22230, USA; Department of Biophysics, Johns Hopkins University, Baltimore, MD 21218, USA; Institute of Molecular Biology, Bulgarian Academy of Sciences, 21, G. Bontchev Str., Sofia 1113, Bulgaria; Institut Curie, PSL Research University, Sorbonne Université, CNRS UMR3664, Laboratoire Dynamique du Noyau, 75005 Paris, France

## Abstract

**Background:** We applied 11 mathematical models of tumor growth to clinical trial data available from public and private sources. We have previously described the remarkable capacity for a simple biexponential model of tumor growth to fit thousands of datasets, and to correlate with overall survival. The goal of this study was to extend our analysis to additional tumor types and to evaluate whether alternate growth models could describe the time course of disease burden in the small subset of patients in whom the biexponential model failed.

**Methods:** For this analysis, we obtained data for tumor burden from 17,140 patients with six different tumor types. Imaging data and serum levels of tumor markers were available for 3,346 and 13,794 patients, respectively. Data from patients were first analyzed using the biexponential model to determine rates of tumor growth (***g***) and regression (***d***); for those whose data could not be described by this model, fit of their data was assessed using seven alternative models. The model that minimized the Akaike Information Criterion was selected as the best fit. Using the model that best fit an individual patient’s data, we estimated the rates of growth (***g***) and regression (***d***) of disease burden over time. The rates of tumor growth (***g***) were examined for association with a traditional endpoint (overall survival).

**Findings:** For each model, the number of patient datasets that fit the model were obtained. As we have previously reported, data from most patients fit a simple model of exponential growth and exponential regression (86%). Data from another 7% of patients fit an alternative model, including 3% fitting to a model of constant or linear regression and exponential growth of tumor on the surface and 3% fitting to model of exponential decay on tumor surface with asymmetric growth. As previously reported, we found that growth rate correlates well with overall survival, remarkably even when data from various histologies are considered together. For patients with multiple timepoints of tumor measurement, the growth rate could often be estimated even during the phase when only net regression of tumor quantity could be discerned.

**Interpretation:** The validation of a simple mathematical model across different cancers and its application to existing clinical data allowed estimation of the rate of growth of a treatment resistant subpopulation of cancer cells. The quantification of available clinical data using the growth rate of tumors in individual patients and its strong correlation with overall survival makes the growth rate an excellent marker of the efficacy of therapy specific to the individual patient.

## INTRODUCTION

In assessing the efficacy of new treatments compared to existing options, phase III clinical trials in cancer often rely on progression-free and overall survival as endpoints. Progression is evaluated by obtaining serial radiographic measurements and/or a specific biomarker reflecting tumor quantity such as prostate specific antigen (PSA) or CA19-9 in patients with prostate or pancreatic cancer, respectively. Using RECIST criteria, progression is scored when the quantity of tumor reaches a value 20% above the start or the nadir achieved during treatment. Overall survival is scored relative to the start of protocol treatment. In most cases, the treatment of advanced metastatic cancers proceeds through an initial phase in which the quantity of tumor falls, and after reaching a nadir in tumor burden, proceeds to a phase where the quantity of tumor increases. Best response reflects a fractional percent below baseline at nadir and is scored as a categorical variable. These metrics – PFS, OS, and response rate – offer insight into the magnitude of response but do not quantify rates of growth or regression. This is a key limitation of current assessment metrics because the rate of disease progression is based upon an individual patient’s cancer biology; having a metric of the rate of progression that can be compared among patients with the same disease and on the same treatment can be invaluable in medical decision-making. Such a metric can also be invaluable in assessing the efficacy of different treatments using data from clinical trials obtained from investigators or sponsors or from publicly available data warehouses such as that of the non-profit organization Project Data Sphere LLC (Cary, NC, USA).

Using data collected from patients with multiple tumor types, we previously estimated the rates of tumor regression and growth using a straightforward model of exponential growth and regression and found that the data in the majority of cases fit well to this model, allowing one to estimate rate constants for tumor growth (***g***) and regression (***d***) that correlate with overall and progression-free survival (Stein et al., 2012; Blagoev et al., 2013; Blagoev et al., 2014; Wilkerson et al., 2017; Leuva et al., 2019; Maitland et al., 2020; Dromain et al., 2021; Yeh et al. 2022). In the present study, in addition to the validated exponential regression and growth models we set out to evaluate an additional seven mathematical models of tumor regression and growth, describing different scenarios including the killing (regression) of cells on the surface or on the surface as well as in the bulk of the tumor; and growth on the surface or on the surface as well as in the bulk of the tumor and asked the question whether any of these models could describe an individual patient’s tumor burden time course when the data failed to fit the simple exponential models. We analyzed tumor quantity data from thousands of patients with solid tumors, estimating the rates of growth (***g***) and regression (***d***) of the fractions of tumor resistant and sensitive to treatment, respectively, for each individual patient.

## METHODS

### Study design, data, and participants

We collected de-identified patient level data from several sources: (1) Project Data Sphere, LLC (PDS), an independent initiative of the CEO Roundtable on Cancer’s Life Sciences Consortium (www.ProjectDataSphere.org); (2) randomized phase III studies sponsored by the pharmaceutical industry who also provided the data (Bristol Myers Squibb, Pfizer) or who had provided the data to the Yale University Open Data Access (YODA) (Johnson & Johnson); (3) clinical trials conducted by the Eastern Cooperative Oncology Group (ECOG) or in the intramural program of the National Cancer Institute; (4) data from the Celgene CA 046 clinical trial accessed from the Vivli platform; and (4) published real world clinical data including that sourced from the Veterans Administration Informatics and Computing Infrastructure (VINCI, Leuva et al, 2019; Sigel et al, 2021). Six tumor types were studied. Specific clinical trial information can be found in **Supplementary Table 1**.

### Procedures and data analysis

In our previous studies, we used a simple exponential regression/exponential growth model to analyze the kinetics of tumor response to therapy, describing regression/decay (***d***) rates and growth (***g***) rates. **Figure 1A** shows a general exponential decay-growth model, with both components detected (the ***gd*** model). The parameter, φ, represents the initial fraction of tumor that is treatment sensitive and 1-φ the initial fraction resistant to treatment. If only decay (tumor regression) can be detected, the biexponential model reduces to a single decaying exponential (φ = 1) and is described as the dx model; if only growth can be fit, the biexponential model reduces to a single growing exponential (φ = 0) and is described as the gx model. In some cancers, such as in kidney cancer, ***g*** and ***d*** are constant during treatment (Burotto et al, 2014). In others, such as in pancreatic cancer, an increase in ***g*** is observed as treatment begins to fail (Yeh et al, 2023). The exponential models describe tumor cells dividing in the whole volume of the tumor, and volumetric data such as that derived from specialized radiomic measurements, and also represented by tumor marker measurements fit very well to the equations (Maitland et al., 2020)

**Figure 1.**
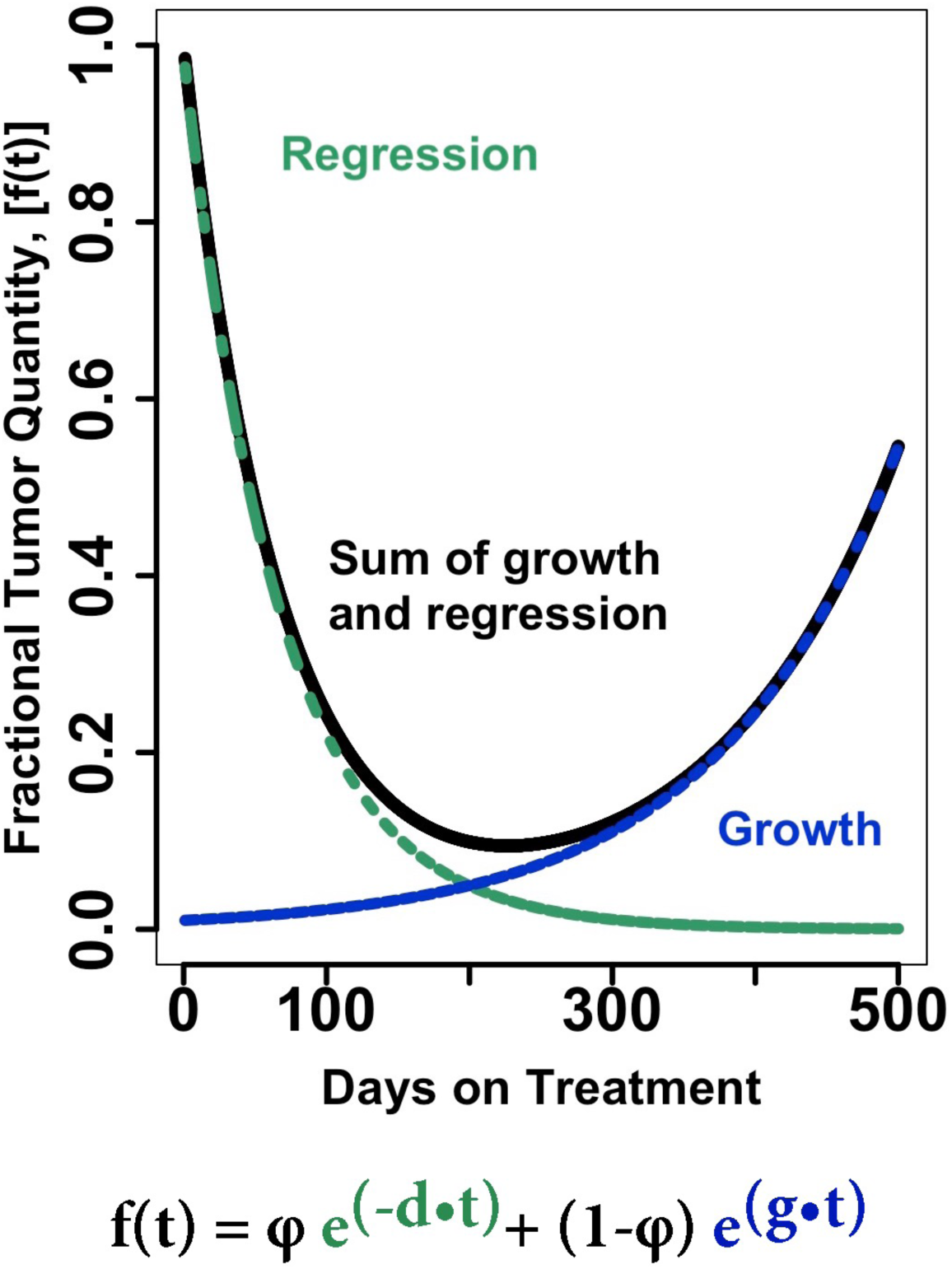
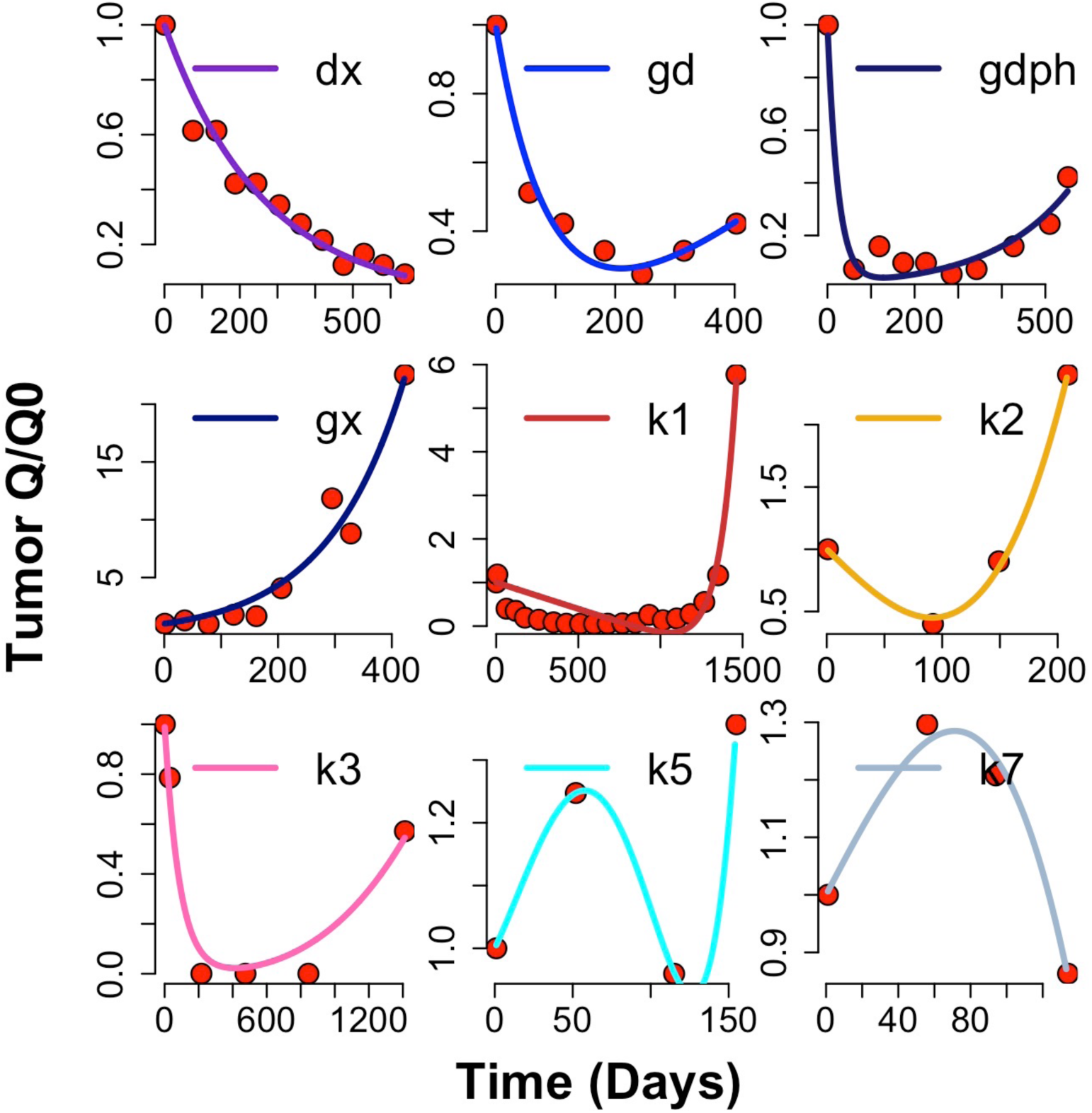
[**A**] **Model for regression and growth**, where f is the fraction or quantity of tumor at time t; φ is the fraction of tumor that is sensitive to the administered therapy and decays or regresses exponentially at rate d; 1– φ the fraction of tumor resistant to the administered therapy and ***g*** the rate of growth exponentially of the resistant fraction. [**B**] **Examples of curve fits**, showing examples of nine of the models tested

However, in a small fraction of patients, typically 5 – 15%, tumor measurement data do not fit well to the equations. Most commonly this is due to inadequate data or data-entry errors, but it is possible that in some cases the dividing cells are mostly on the tumor surface and/or that drug action is restricted to the surface due to poor penetration, as proposed for pancreatic cancer (Erkan et al., 2012). To assess these possibilities, we developed mathematical models where tumor growth and/or regression occur only on the surface or in the bulk of the tumor (**Table 1**). In addition to modeling treatments that kill a ***constant fraction*** of all tumor cells, leading to an exponential tumor decay, we also modeled a treatment that kills a ***constant number*** of cells with each drug administration, leading to a linear tumor decay. Modelling tumor growth included growth on the surface only, or on the surface and in the bulk of the tumor. We also included growth through ***symmetric*** or ***asymmetric*** cancer cell divisions. In an ***asymmetric*** cell division model, one of the offspring has finite proliferative potential and cannot replicate indefinitely, but the other can, leading to linear tumor growth; while in a ***symmetric*** cell division model, both offspring can divide, leading to an exponential growth when the cells are growing in the whole tumor (Blagoev et al., 2021). The mathematical derivation of the equations describing these different decay and growth patterns are described in Supplementary Materials. We asked whether equations described by any of these models could be fit to the data that were not explained by the simpler biexponential model. **Figure 1B** includes examples of curve fits for nine of the 11 models. While two of the alternative models demonstrate fits of the data comprising 3% of patients each, the remaining alternative models that were fit by the algorithm show data that almost certainly represent artifact.

**Table 1.** Summary of exponential and alternative mathematical models used in the analysis. The four exponential models assume the regression process follows exponential decay in the bulk and the growth process follows exponential growth in the bulk. The seven alternative models explore alternative assumptions about the two processes. The mathematical equations used are shown with ***g*** representing the growth rate constant, ***d*** representing the decay/regression rate constant and **φ** representing the fraction of tumor that is sensitive to the therapy.

| Table 1 – Summary of exponential and alternative mathematical models used in the analysis |  |  |  |
| --- | --- | --- | --- |
| Exponential Models |  |  |  |
| Mechanism |  | Model | Equation |
| Regression only | | dx | $f(t) = e^{-dt}$ |
| Growth only | | gx | $f(t) = e^{gt}$ |
| Both regression and growth | | gd | $f(t) = e^{-dt} + e^{gt} - 1$ |
| Both regression and growth with fraction sensitive | | gdphi | $f(t) = \phi e^{-dt} + (1 - \phi)e^{gt}$ |
| Alternative Models |  |  |  |
| Regression Mechanism | Growing Mechanism | Model | Equation |
| Constant Number | Exponential growth, Bulk | k1 | $f(t) = (\phi - dt) + (1 - \phi)e^{gt}$ |
| Constant Number | Exponential growth, Surface | k2 | $f(t) = (1 - dt) + gt^3$ |
| Exponential decay, Bulk | Exponential growth, Surface | k3 | $f(t) = \phi e^{-dt} + 1 - \phi + gt^3$ |
| Exponential decay, Bulk | Asymmetric | k4 | $f(t) = \phi e^{-dt} + 1 - \phi + gt$ |
| Exponential decay, Surface | Exponential growth, Bulk | k5 | $f(t) = (\phi - dt^3) + (1 - \phi)e^{gt}$ |
| Exponential decay, Surface | Exponential growth, Surface | k6 | $f(t) = (1 - dt^3) + gt^3$ |
| Exponential decay, Surface | Asymmetric | k7 | $f(t) = (1 - dt^3) + gt$ |

## RESULTS

For this analysis, we obtained data of tumor burden from 22,383 patients and analyzed the data from 17,140 who had at least three tumor measurements. Imaging data and serum levels of tumor markers were available for 3,346 and 13,794 patients, respectively. Radiographic imaging was available for patients with a diagnosis of breast, colorectal and non-small cell lung (NSCLC) cancers and in these we used the sum of longest diameter according to RECIST as a measure of tumor burden. Tumor markers included PSA values in patients with D0 and castration-resistant prostate cancer (CRPC), M-spikes in multiple myeloma (MM) and its precursor states and CA19-9 in pancreatic cancer. **Supplemental Table 1** summarizes the types of cancers and the treatments received for those whose data were used in this study. Note that the data in all cases was obtained as patients received treatment, a majority while enrolled in a clinical trial and others while receiving treatment in real-world settings. Data from the 17,140 patients were initially analyzed using the four exponential models in **Table 1** (Stein et al., 2012; Blagoev et al., 2013; Blagoev et al., 2014; Wilkerson et al., 2017; Leuva et al., 2019; Maitland et al., 2020; Dromain et al., 2021; Yeh et al. 2022) followed by the seven alternative models for those data not fitting the simple exponential model. The model that optimally minimized the Akaike Information Criterion (AIC) was selected as the best fit. The corresponding ***g***, ***d***, and φ values were extracted from the model selected as best fitting data. Percentages of each model fit and the medians of the growth (***g***) and regression (***d***) rates for each model are shown. The number of patients that fit each particular model are shown in **Table 2**. **Panel A** summarizes the results for the entire data set while **Panels B** and **C** report the data according to disease category. Note how for all histologies, the percentage of the data that fit the ***dx*** and ***gx*** models, is higher and lower, respectively, during treatments used in first line versus treatments administered in the second line setting, reflecting the evolution to a less drug sensitive tumor. The alternative models of tumor growth consider variables relating to growth/regression on surface, asymmetric growth, and constant vs. exponential growth/regression. Only a small number of patient datasets were well-described by these alternative models, namely 3% of data fitted model k2 and 3% fit model k7 (see **Table 1**). The ***g*** values derived from model fits to the k2 and k7 equations were not included in subsequent Kaplan-Meier curves.

**Table 2.**
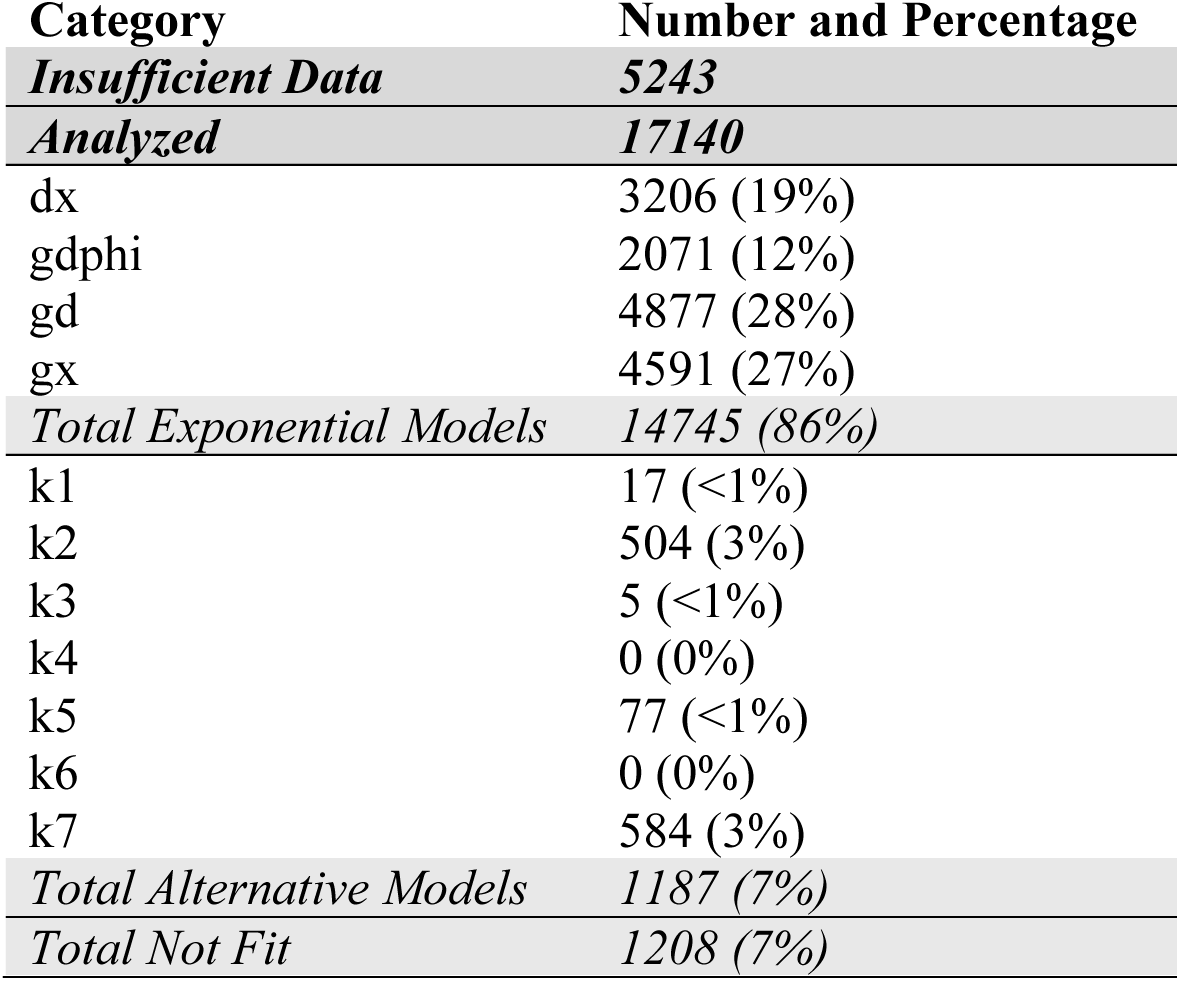
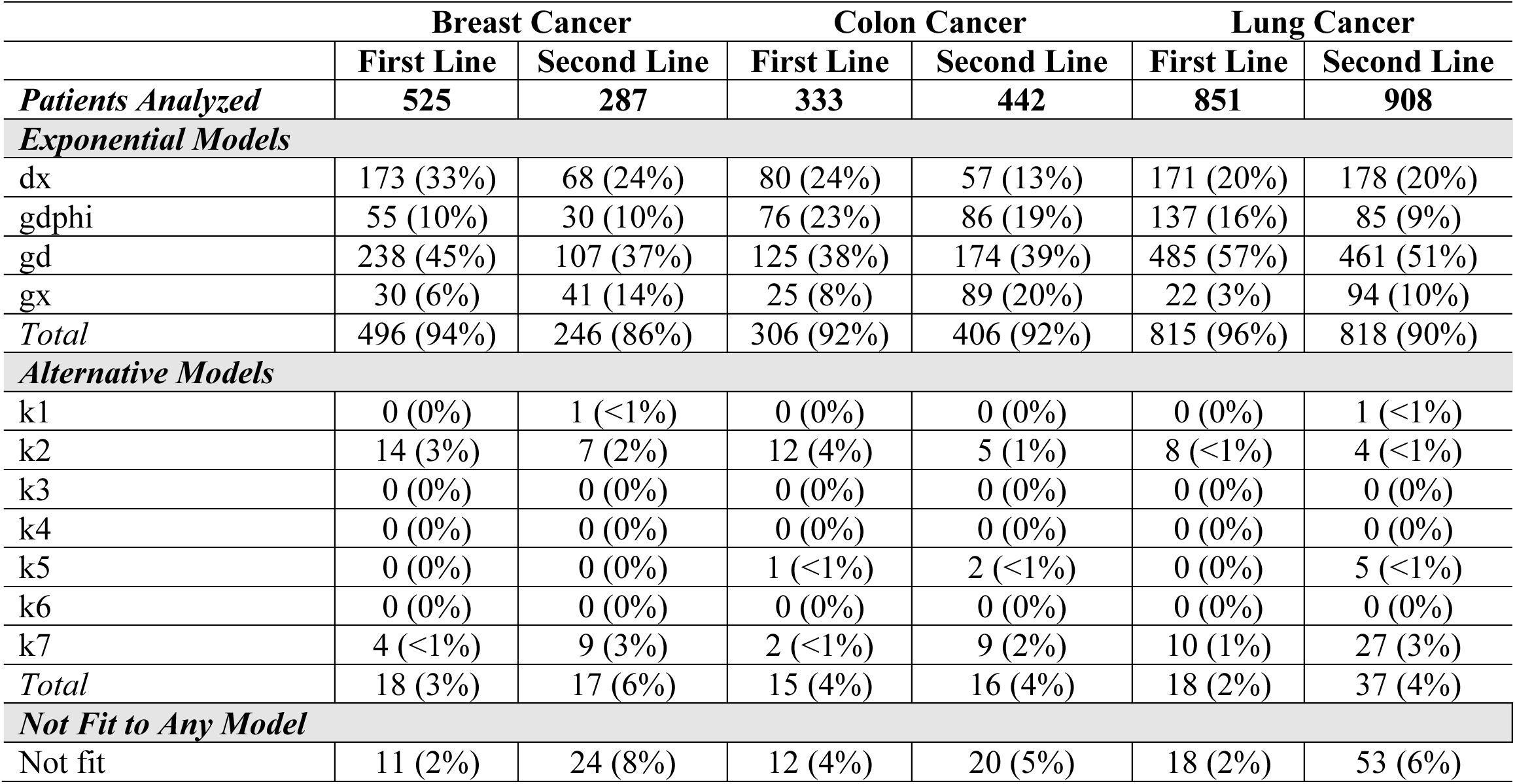
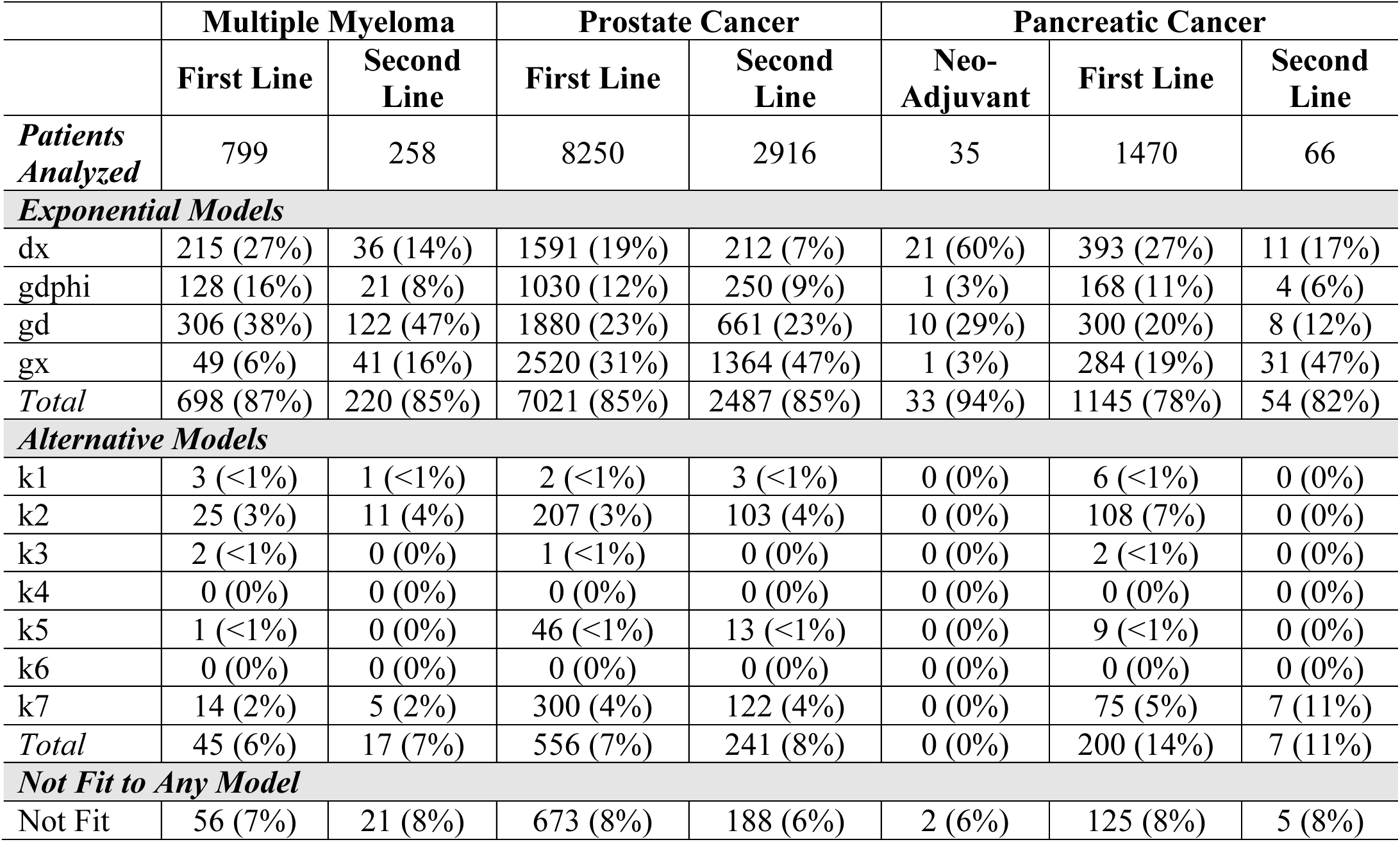
Summary of data analysis and models by type of tumors and line of therapy: The data from patients with at least three imaging or serum biomarker measurements (N=17,140) were analyzed with the four exponential models and if not fit to any of the models, analyzed with the seven alternative models. **Panel A – Model fits, all patients. Panel B – Model fits, according to disease category; tumor burden measured by radiographic imaging. Panel C – Model fits, according to disease category; tumor burden measured by serum biomarkers.**

| Table 2, Panel A – Model fits in all patients |  |
| --- | --- |
| Category | Number and Percentage |
| *Insufficient Data* | *5243* |
| *Analyzed* | *17140* |
| dx | 3206 (19%) |
| gdphi | 2071 (12%) |
| gd | 4877 (28%) |
| gx | 4591 (27%) |
| *Total Exponential Models* | *14745 (86%)* |
| k1 | 17 (<1%) |
| k2 | 504 (3%) |
| k3 | 5 (<1%) |
| k4 | 0 (0%) |
| k5 | 77 (<1%) |
| k6 | 0 (0%) |
| k7 | 584 (3%) |
| *Total Alternative Models* | *1187 (7%)* |
| *Total Not Fit* | *1208 (7%)* |

**Table 2, Panel B – Disease-specific fits, tumor burden measured by radiographic imaging**
|  | **Breast Cancer** |  | **Colon Cancer** |  | **Lung Cancer** |  |
| --- | --- | --- | --- | --- | --- | --- |
|  | **First Line** | **Second Line** | **First Line** | **Second Line** | **First Line** | **Second Line** |
| ***Patients Analyzed*** | **525** | **287** | **333** | **442** | **851** | **908** |
| ***Exponential Models*** |  |  |  |  |  |  |
| dx | 173 (33%) | 68 (24%) | 80 (24%) | 57 (13%) | 171 (20%) | 178 (20%) |
| gdphi | 55 (10%) | 30 (10%) | 76 (23%) | 86 (19%) | 137 (16%) | 85 (9%) |
| gd | 238 (45%) | 107 (37%) | 125 (38%) | 174 (39%) | 485 (57%) | 461 (51%) |
| gx | 30 (6%) | 41 (14%) | 25 (8%) | 89 (20%) | 22 (3%) | 94 (10%) |
| *Total* | 496 (94%) | 246 (86%) | 306 (92%) | 406 (92%) | 815 (96%) | 818 (90%) |
| ***Alternative Models*** |  |  |  |  |  |  |
| k1 | 0 (0%) | 1 (<1%) | 0 (0%) | 0 (0%) | 0 (0%) | 1 (<1%) |
| k2 | 14 (3%) | 7 (2%) | 12 (4%) | 5 (1%) | 8 (<1%) | 4 (<1%) |
| k3 | 0 (0%) | 0 (0%) | 0 (0%) | 0 (0%) | 0 (0%) | 0 (0%) |
| k4 | 0 (0%) | 0 (0%) | 0 (0%) | 0 (0%) | 0 (0%) | 0 (0%) |
| k5 | 0 (0%) | 0 (0%) | 1 (<1%) | 2 (<1%) | 0 (0%) | 5 (<1%) |
| k6 | 0 (0%) | 0 (0%) | 0 (0%) | 0 (0%) | 0 (0%) | 0 (0%) |
| k7 | 4 (<1%) | 9 (3%) | 2 (<1%) | 9 (2%) | 10 (1%) | 27 (3%) |
| *Total* | 18 (3%) | 17 (6%) | 15 (4%) | 16 (4%) | 18 (2%) | 37 (4%) |
| ***Not Fit to Any Model*** |  |  |  |  |  |  |
| Not fit | 11 (2%) | 24 (8%) | 12 (4%) | 20 (5%) | 18 (2%) | 53 (6%) |

|  | **Multiple Myeloma** |  | **Prostate Cancer** |  | **Pancreatic Cancer** |  |  |
| --- | --- | --- | --- | --- | --- | --- | --- |
|  | **First Line** | **Second Line** | **First Line** | **Second Line** | **Neo-Adjuvant** | **First Line** | **Second Line** |
| ***Patients Analyzed*** | 799 | 258 | 8250 | 2916 | 35 | 1470 | 66 |
| ***Exponential Models*** |  |  |  |  |  |  |  |
| dx | 215 (27%) | 36 (14%) | 1591 (19%) | 212 (7%) | 21 (60%) | 393 (27%) | 11 (17%) |
| gdphi | 128 (16%) | 21 (8%) | 1030 (12%) | 250 (9%) | 1 (3%) | 168 (11%) | 4 (6%) |
| gd | 306 (38%) | 122 (47%) | 1880 (23%) | 661 (23%) | 10 (29%) | 300 (20%) | 8 (12%) |
| gx | 49 (6%) | 41 (16%) | 2520 (31%) | 1364 (47%) | 1 (3%) | 284 (19%) | 31 (47%) |
| ***Total*** | 698 (87%) | 220 (85%) | 7021 (85%) | 2487 (85%) | 33 (94%) | 1145 (78%) | 54 (82%) |
| ***Alternative Models*** |  |  |  |  |  |  |  |
| k1 | 3 (<1%) | 1 (<1%) | 2 (<1%) | 3 (<1%) | 0 (0%) | 6 (<1%) | 0 (0%) |
| k2 | 25 (3%) | 11 (4%) | 207 (3%) | 103 (4%) | 0 (0%) | 108 (7%) | 0 (0%) |
| k3 | 2 (<1%) | 0 (0%) | 1 (<1%) | 0 (0%) | 0 (0%) | 2 (<1%) | 0 (0%) |
| k4 | 0 (0%) | 0 (0%) | 0 (0%) | 0 (0%) | 0 (0%) | 0 (0%) | 0 (0%) |
| k5 | 1 (<1%) | 0 (0%) | 46 (<1%) | 13 (<1%) | 0 (0%) | 9 (<1%) | 0 (0%) |
| k6 | 0 (0%) | 0 (0%) | 0 (0%) | 0 (0%) | 0 (0%) | 0 (0%) | 0 (0%) |
| k7 | 14 (2%) | 5 (2%) | 300 (4%) | 122 (4%) | 0 (0%) | 75 (5%) | 7 (11%) |
| ***Total*** | 45 (6%) | 17 (7%) | 556 (7%) | 241 (8%) | 0 (0%) | 200 (14%) | 7 (11%) |
| ***Not Fit to Any Model*** |  |  |  |  |  |  |  |
| Not Fit | 56 (7%) | 21 (8%) | 673 (8%) | 188 (6%) | 2 (6%) | 125 (8%) | 5 (8%) |

**Table 3.** Overall survivals calculated using the estimated *g* values and regression equation in Figure 3 and OS survivals reported in the literature.

| **Disease** | **Cohort** | **Treatment** | **Median *g* value** | **Estimated median OS using equation OS = 0.0077 x (1/*g*) + 12.52** | **Median OS from published report** |
| --- | --- | --- | --- | --- | --- |
| **Breast Cancer** | PALOMA 1 | Palbociclib + Letrozole | 0.000268 | 41.4 | 37.5 |
|  |  | Letrozole | 0.000468 | 29.0 | 34.5 |
|  | PALOMA 2 | Palbociclib + Letrozole | 0.000136 | 69.3 | 53.9 |
|  |  | Letrozole | 0.000137 | 68.9 | 51.2 |
| **Colon Cancer** | Amgen PRIME | Panitumumab + FOLFOX | 0.000877 | 21.3 | 23.9 |
|  |  | FOLFOX | 0.001318 | 18.4 | 19.7 |
| **Lung Cancer** | AstraZe_2006_152 | Paclitaxel + Carboplatin | 0.001479 | 17.7 | 18.3 |
|  | Celgene_2007_108 | Paclitaxel + Carboplatin | 0.001392 | 18.1 | 11.2 |

As previously reported in prostate, colorectal, and pancreatic cancer (Leuva et al, 2019; Maitland et al, 2020; Yeh et al, 2023), and observed by us in all cancers, the growth rates in the patients analyzed in this study correlate very well with overall survival (OS) as shown in **Figure 2**. In this plot, we split the data into quantiles of growth rates and show that the overall survival of each quantile correlates inversely with the growth rate. While it is expected that how fast tumors grow, or how aggressive they are would be related with the time to death, it is remarkable that this relationship transcends tumor type as shown in panel A, and that the growth rates measured over a brief fixed period of time, as during a clinical trial, predict a survival date that occurs in most cases much later in time, often years later. We also assessed the mathematical relationship between the OS and the growth rate. We found that as one would intuitively predict, OS should be inversely proportional to the growth rate, as shown in **Figure 3**.

**Figure 2.**
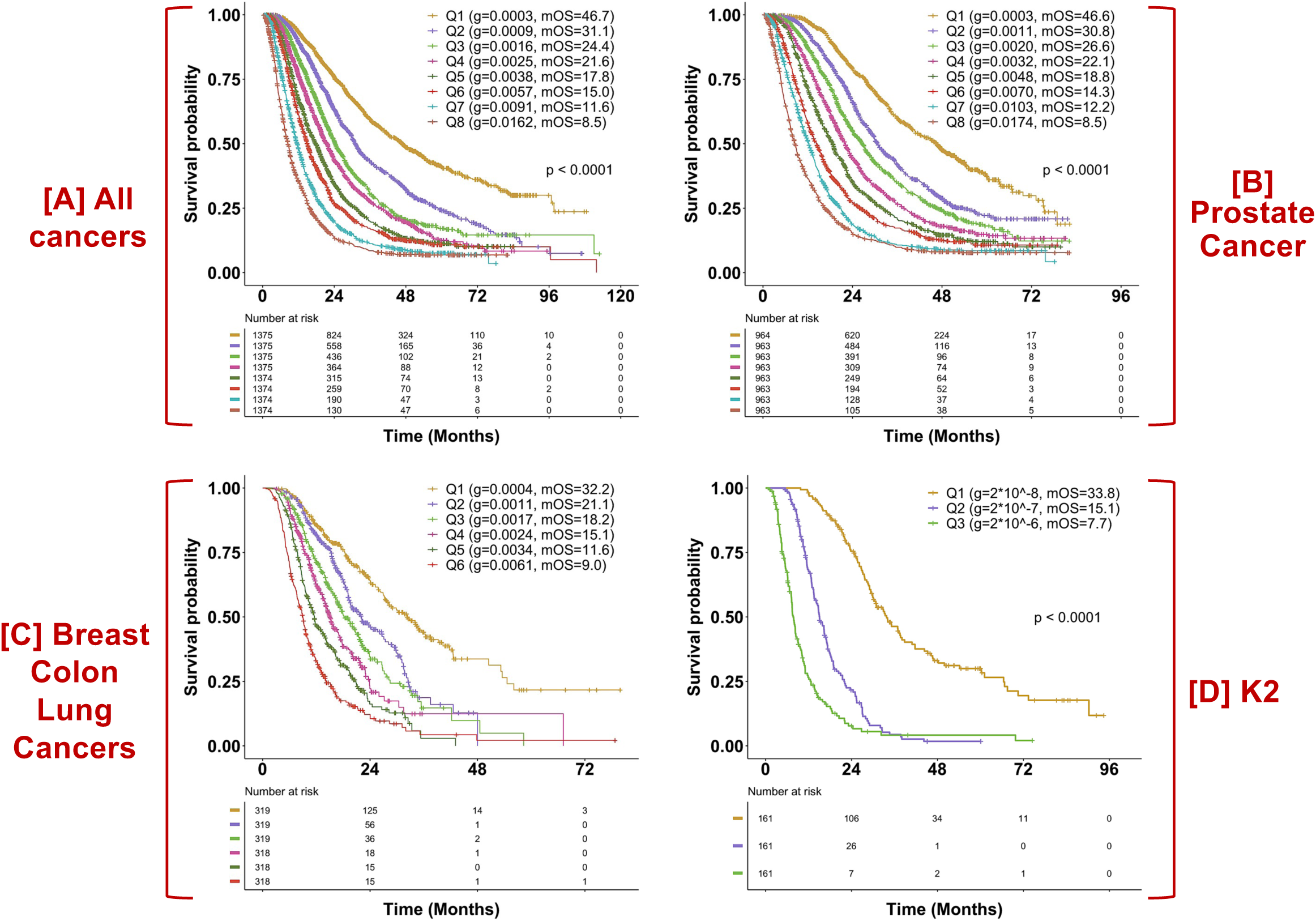
Correlation with overall survival (OS) Kaplan-Meier (KM) plots of OS by quantiles of the growth rate constant ***g*** are displayed in [**A**] octiles all patients [**B**] octiles of patients diagnosed with prostate cancer [**C**] quartiles of patients diagnosed with breast, colon and lung cancers [**D**] terciles of patients whose data gave a best fit of k2. In each case, the data was divided into quantiles according to the ***g*** values and then for each quantile a KM of OS was plotted. Log-rank test was used to assess the differences between curves.

**Figure 3.**
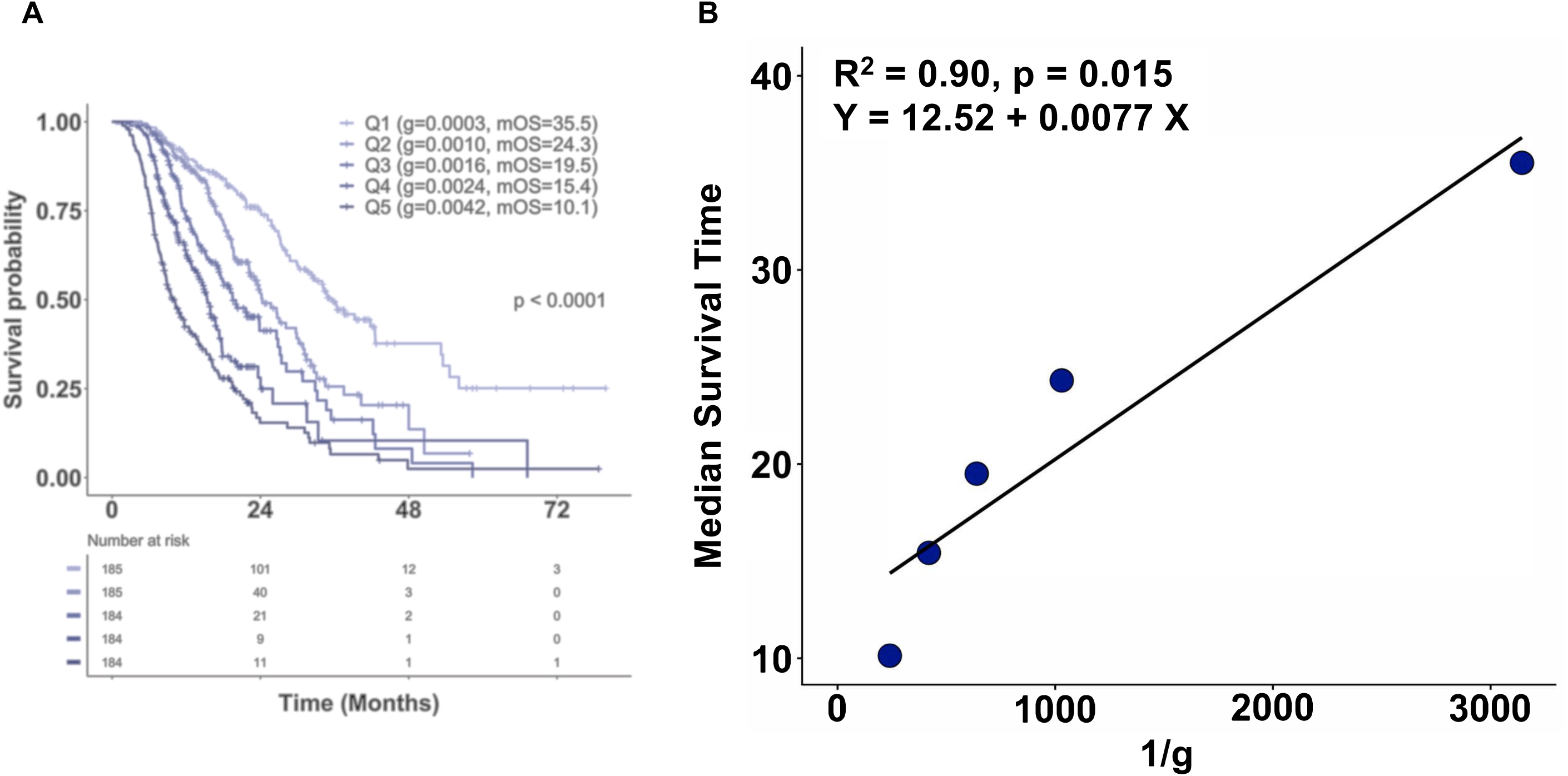
Correlation of overall survival and *g* values. [**A**] Kaplan-Meier (KM) plots of OS by quintiles of the growth rate constant ***g*** of patients diagnosed with breast, colorectal and lung cancers receiving a therapy administered in first line. As in Figure 2, the data was divided into quintiles according to the ***g*** values and then for each quintile a KM of OS was plotted. Log-rank test was used to assess the differences between curves. [**B**] For each of the quintiles, the median ***g*** values and median survival time (months) extracted from the Kaplan-Meier plot in panel [A] were plotted. The black dots represent the actual values of median survival time plotted on the Y-axis and 1/median ***g*** values on the X-axis.

An important question a patient and their doctors have during treatment is how well is the treatment working? Often this question cannot be answered early on in the treatment and even when data on progression of the disease are available, quantitative measures of progression are rare outside of a clinical trial. Our discovery that the growth rate obtained from our simple mathematical model correlates very well with OS, gives patients and the medical professionals a tool to compare the growth rate of the tumor in a patient with a cohort of patients with the same cancer and treatment. A distribution of growth rates of tumors in patients categorized as having complete response (CR), partial response (PR), stable disease (SD), and progressive disease (PD) is shown in **Figure 4**.

**Figure 4.**
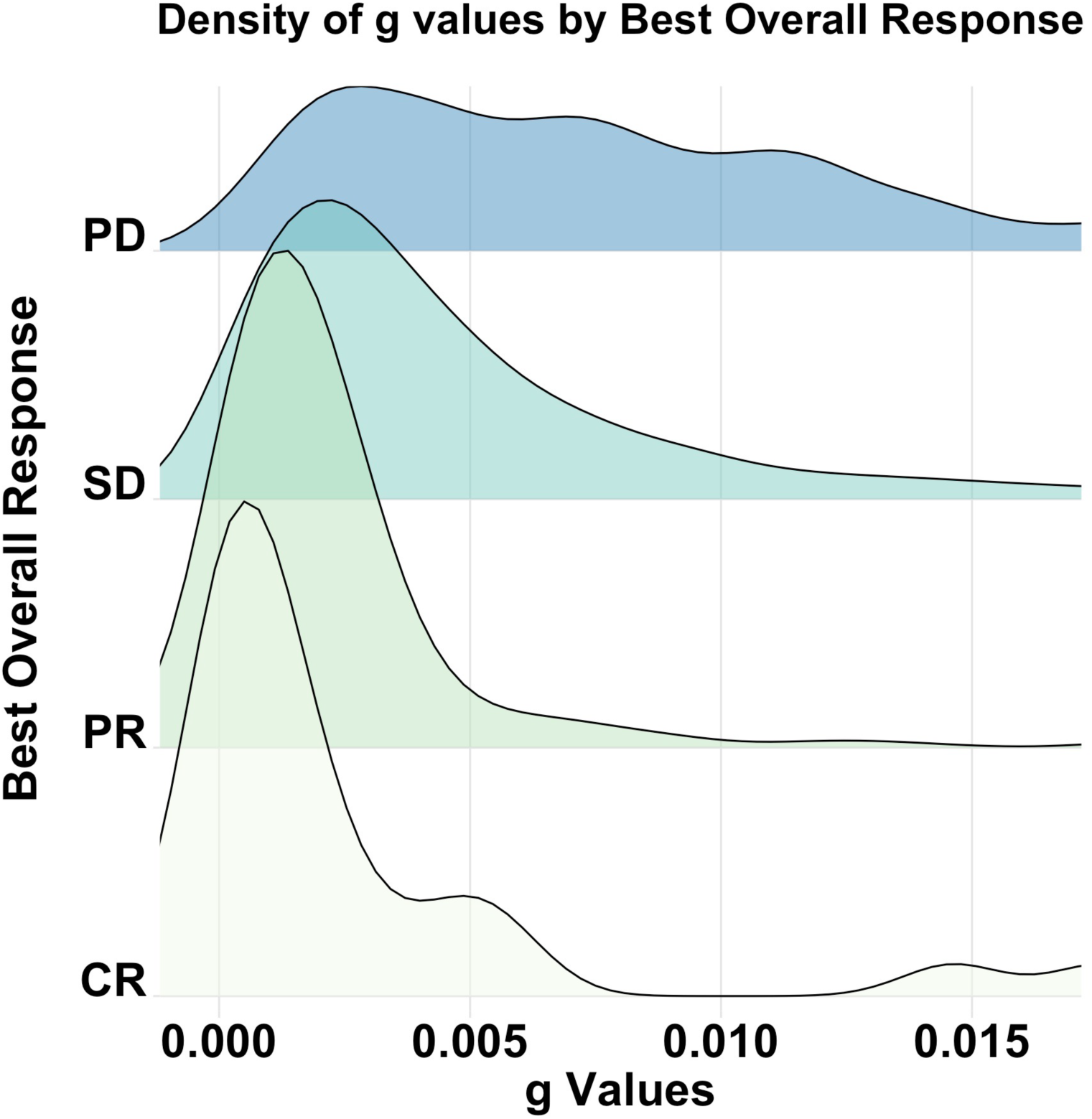
Distribution of *g* values in patients according to best RECIST response. Best overall response of complete response (CR), partial response (PR), stable disease (SD) and progressive disease (PD).

In addition to the significant correlation with OS, the growth rate also correlates with progression-free survival (PFS) as shown in **Figure 5A**. Time to progression is the time from randomization to tumor progression. PFS is a similar endpoint, but also includes death from all causes. These correlations show that the methodology can be used not only for evaluating individual patient’s treatments, but also can be informative in clinical trials.

**Figure 5.**
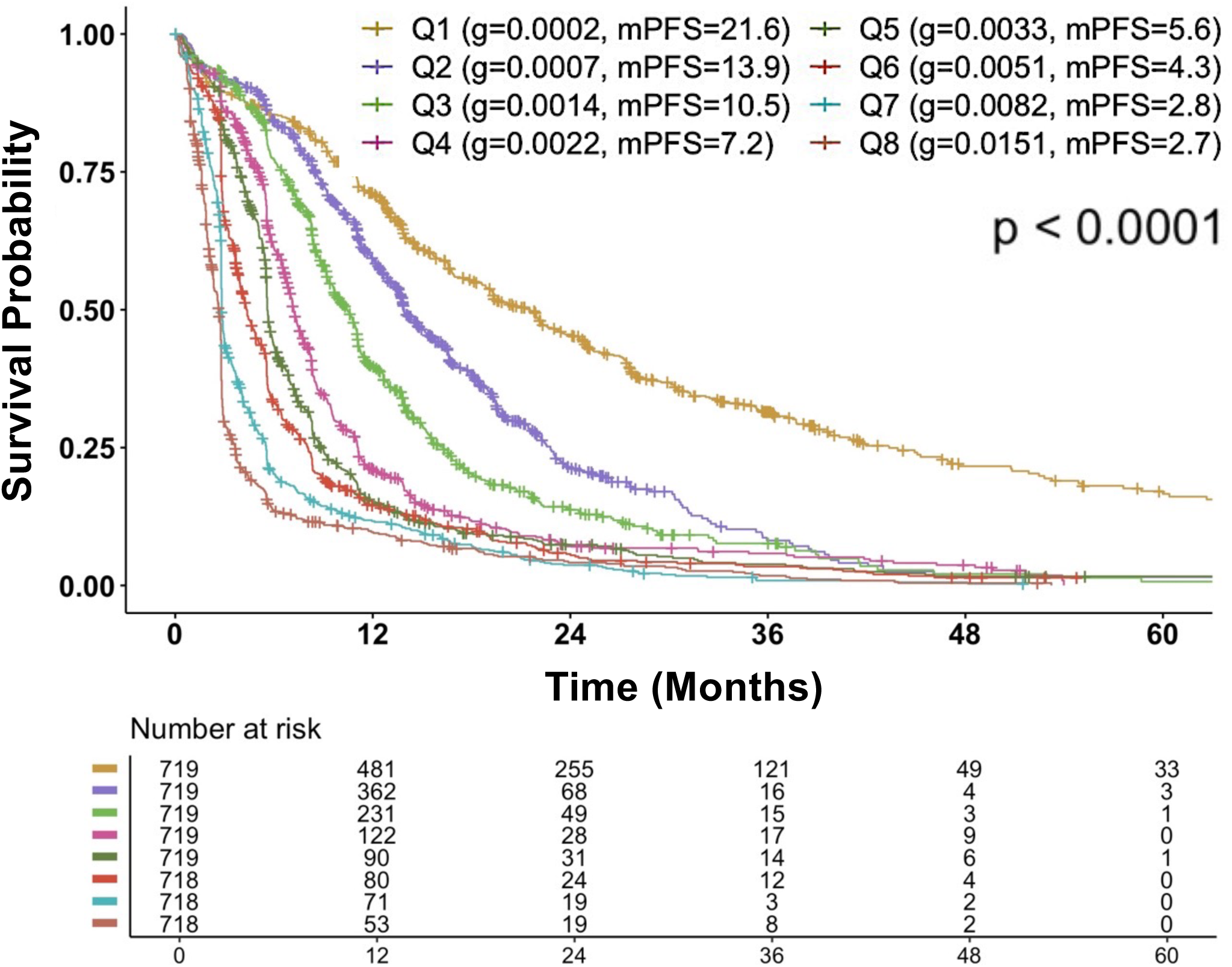
Correlation with progression-free survival. Kaplan-Meier plots of PFS by octile of the growth rate constant g are displayed. Log-rank test was used to assess the differences between curves.

## DISCUSSION

In this analysis we have used parametric machine learning to analyze the kinetics of a large number of tumors representing several different cancer types. While we used many models of tumor growth, we found that the vast majority of patients’ data fit to a model in which tumor kinetics are well-captured by the sum of concurrent exponential rates of decay and a growth. Each of these rates reflects the subset of cells sensitive to the treatment (decay) and the subset of cells insensitive to it (growth), respectively. From our analysis it is unclear whether the treatment induces resistance or simply selects for resistant clones; the initial stability of the rate of the growing fraction suggests that treatment allows the resistant fraction to emerge. However, in some tumor types, such as pancreatic cancer, a faster rate of growth is observed at the end of treatment than during the main period of treatment, suggesting resistance may emerge during treatment.

Another purpose of this study was to examine alternative models of tumor growth, by considering variations relating to growth and regression on the surface vs. in the total tumor bulk, asymmetric vs. symmetric growth, and constant vs. exponential growth and regression. Only a small number of patient data was better described by other of the models we used, namely 3% of data fitted model k2 and 3% fitted model k7 (see **Table 1**). The first of these models describes cell death that is linear, meaning mean that the treatment kills a constant number of cells per day. The growth part of the model describes cells dividing symmetrically, but only on the surface of the tumor. While 3% of patients had tumor data described by this model, the growth rate, ***g***, is 0.00000002, a rate that translates to a tumor doubling time of 94,931 years! This growth rate is too low to be biologically meaningful and we believe that the fits were caused by noise in the data. In addition, drug action whereby each time a drug administered always kills the same number of cells also seems to be biologically meaningless. The only other model, k7, describes exponential decay of the sensitive cells and linear growth of the resistant cells. In these cases, the linear tumor growth would be the result of asymmetric cancer cell divisions. Although this represented only 584 datasets among those of 17,140 patients, we cannot exclude the possibility these patients had tumors that were biologically more indolent or less aggressive than the vast majority of patients with tumors whose data was best described by exponential growth. Alternatively, some investigators have published growth models derived from cancer stem cells, in which growth was asymmetric. The k7 model may have captured a population of patients in which this was measurable.

We also examined correlations of ***g*** with overall survival. The high correlations of an event such as survival often delayed by years from the time when patients were receiving a treatment and tumor burden determined and ***g*** values estimated using data captured only while the patient is enrolled in a clinical trial suggests that both the impact of the therapy as well as an inherent biology of the tumor are reflected in ***g*** values This should not be surprising given that more aggressive tumors that grow faster also often respond poorly to therapeutic interventions. Correlations with overall survival could be seen even when data from a wide range of cancers were blended together. While unexpected, this may in fact be foreseeable given this data comes from patients with metastatic cancers whose disease shares aggressiveness with other metastatic cancers and whose overall survival is often not too dissimilar, given the limitations of our therapies and their inability in most cases to cure cancer. Additionally, we show in this study that the distribution (density plot) of ***g*** values reflects RECIST-designated response definitions. It is interesting to note that patients designated with disease progression as best response have the broadest distribution of ***g*** (**Figure 4**), and that effective therapy as seen in PR and CR is associated with a much narrower range of distribution. One implication of this type of analysis is that an individual patient’s ***g*** value could be overlaid on the density plots to determine whether a patient’s ***g*** is in a satisfactory range, and consistent with a RECIST response metric even prior to that metric having been met.

Finally, we set out to determine how well the ***g*** values could predict OS by using the ***g*** values we estimated and inserting these into the equation in **Figure 3** that describes the regression and comparing that OS with the reported OS. As noted above, the high correlation between ***g*** using data harvested while a patient receives a therapeutic and OS often years later suggests ***g*** “reads out” not only sensitivity to the therapeutic that is being administered but likely also the inherent biology of the tumor. For this example, we used data from breast, colorectal, and pancreatic cancer obtained by imaging receiving a therapy administered in first line. **Figure 6** shows the correlation between the overall survivals predicted using the estimated ***g*** values and the regression equation in **Figure 3** and overall survivals reported in the literature. The high correlation (R^2^=0.9) demonstrates the ability to use ***g*** to predict OS and allows for target ***g*** values to be calculated that can achieve desired improvements in OS.

**Figure 6.**
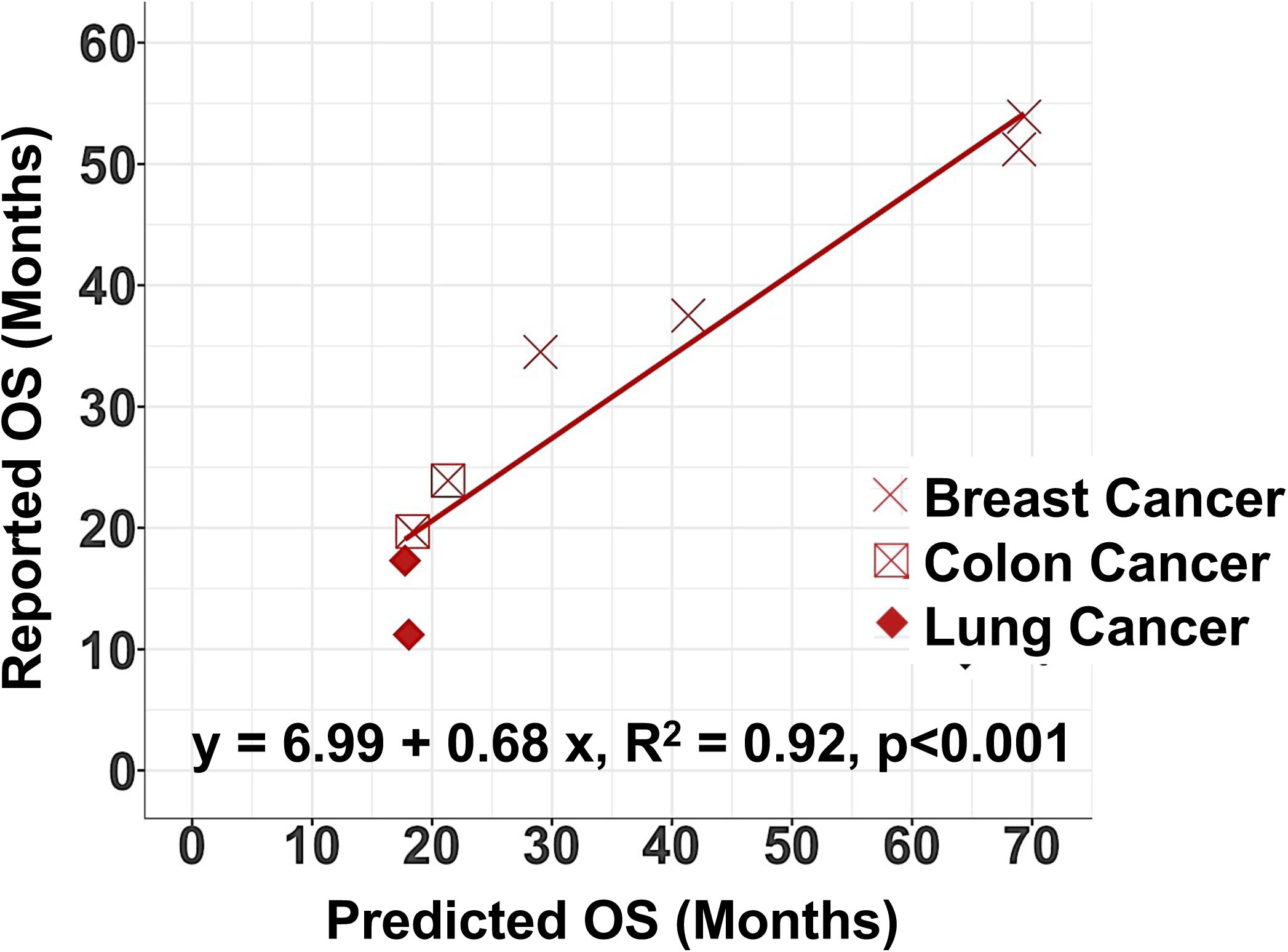
*g* values can be used to predict overall survival. Comparison of overall survivals predicted using the estimated ***g*** values estimated as part of the present study and the equation that describes the regression in Figure 3 and the overall survivals reported in the literature.

In sum, we have extended our prior disease-specific studies to show that rates of tumor growth estimated using data obtained while a patient receives a treatment is inversely correlated with progression-free and overall survival in a way that transcends tumor type. We have tested alternative models for those 14% of tumors in which there are poor fits to the simple biexponential model, and found that only 7% fit an alternative model, and of those, only 3% fit in a biologically meaningful way. We thus consider the biexponential model to be the most biologically and practically sound model of tumor growth available.

## ACKNOWLEDGEMENTS

The National Science Foundation supported the work of K. B. Blagoev. The National Science Foundation had no role in study design, data collection and analysis, decision to publish, or preparation of the manuscript. The views presented here are not those of the National Science Foundation and represent solely the views of the authors.

## Author contributions

KBB, TF, and SEB designed the study, KBB, and MZ analyzed the data. KBB, TF, LS, MZ and SEB wrote the paper.

## Competing interests

The authors declare no competing interests.

## Funding

Krastan Blagoev was funded through the National Science Foundation of USA Independent Research and Development Program. Susan Bates and Tito Fojo are supported in part by the Veterans Health Administration, the Herbert Irving Comprehensive Cancer Center (TF, SB), the Blavatnik Foundation (TF) and the Prostate Cancer Foundation (TF); the Pancreas Center (SB) and the Pancreatic Cancer Action Network (SB).

## Notes

### Competing Interest Statement

The authors have declared no competing interest.

